# Mechanism of Mycolactone Toxin Membrane Permeation: Atomistic vs Coarse-Grained MARTINI Simulations

**DOI:** 10.1101/470807

**Authors:** F. Aydin, R. Sun, J. M. J. Swanson

## Abstract

Mycolactone, a cytotoxic and immunosuppressive macrolide produced by *Mycobacterium ulcerans*, is the central virulent factor in the skin disease Buruli ulcer. This multifunctional cytotoxin affects fundamental cellular processes such as cell adhesion, immune response and cell death by targeting various cellular structures. Developing effective diagnostics that target mycolactone has been challenging, potentially due to suspected interactions with lipophilic architectures, including membranes. To better understand the pathogenesis of Buruli ulcer disease, aid in the development of diagnostics, and learn how amphiphiles in general use lipid trafficking to navigate the host environment, we seek to understand the nature of mycolactone-membrane interactions. Herein we characterize how the two dominant isomers of mycolactone (A and B) interact with and permeate DPPC membranes with all-atom molecular dynamics simulations employing transition tempered metadynamics, and compare these results with those obtained by MARTINI coarse-grained simulations. Our all-atom simulations reveal that both isomers have a strong preference to associate with the membrane, although their mechanisms and energetics of membrane permeation differ slightly. Water molecules are found to play an important role in the permeation process. Although the MARTINI coarse-grained simulations give the correct free energy of membrane association, they fail to capture the mechanism of permeation and role of water during permeation as seen in all-atom simulations.

## INTRODUCTION

Mycolactone is an exotoxin produced by *Mycobacterium ulcerans* and the central causative agent behind the neglected tropical skin disease called Buruli ulcer (1). The main characteristics of Buruli ulcer disease are extensive tissue destruction (due to the cell death) and a surprisingly low level of pain or inflammation (2, 3). The toxin is thought to be secreted in bacterial outer membrane vesicles and then delivered to host cells (4) where it is cytotoxic and immunosuppressive, disrupting various cellular functions such as cytoskeletal organization, cytokine and chemokine expression and other signaling cascades (5, 6). Although multiple cellular targets have been identified, the majority of toxin’s cellular effects can likely be explained by its inhibition of the sec61 translocon (7-9). By binding to the translocon pore the toxin has been shown to block co-translational translocation of secretory proteins, leading to numerous downstream consequences. Other cellular targets include angiotensin II receptors (AT2R) expressed by neurons (10, 11), and the Wiskott-Aldrich syndrome protein (WASP) to which mycolactone binds with a 100-fold higher binding affinity than CDC42, the strongest natural activator of WASP (12). Binding of mycolactone causes WASP to open, exposing a domain that activates the Arp2/3 complex (13), which in turn nucleates branched filamentous actin (F-actin) and results in excessive actin branching. Excessive actin branching is associated with abnormal cell adhesion and uncontrolled cell migration that can also induce to apoptosis of host cells.

It is unknown why the toxin has so many biological targets, or what unifies its pleiotropic effects. One interesting possibility arises from the toxin’s expected interactions with membranes. Both AT2R and WASP are membrane associated proteins that have been suggested to be regulated by ordered microdomains that localize and facilitate assembly of signaling complexes (14-16). Our previous computational work suggested that toxin decreases line tension, thereby disrupting domain formation (17). Recent experimental work confirmed this prediction. Using Lagmuir monolayers to mimic the plasma membrane, mycolactone was shown to preferentially bind membranes with cholesterol and destabilize the lipid ordered phase, disrupting domain formation and fluidizing the membrane. This finding suggests that mycolactone’s influence on membrane reshaping could unify or at least enhance its pleitropic effects. Whether this hypothesis is true or not, mycolactone is clearly a complex toxin that impairs cellular function in multiple ways.

Although treatment regimens exist and are improving (18, 19), early detection of Buruli ulcer disease remains a significant challenge. Current diagnostic strategies include microscopic detection of acid-fast bacilli, cultures, PCR targeting specific M. ulcerans genes, and histopathology (20). However, none of these strategies is rapid, reliable, or field deployable. Targeting the toxin itself in a diagnostic assay would be ideal and has been identified as a central research aim by the World Health Organization (19). Despite numerous efforts toward this goal however, rapid diagnostic assays targeting mycolactone have yet to be identified. One promising advance was the development of mycolactone-specific monoclonal antibodies (21). Unfortunately, these antibodies have only proven to be effective and/or neutralizing if they are pre-equilibrated with the toxin before exposure to host cells. One possible cause of these challenges, which is consistent with the toxin’s amphipathic (or amphiphilic) structure, is that that toxin is hidden from tracking antibodies due to association with lipophilic carriers such as membranes.

In this work, we aim to further characterize the nature of mycolactone-membrane interactions to better understand its mechanism of host cell penetration and distribution in host environment, factors highly relevant to the toxin’s pathogenicity and the design of effective diagnostics. We are additionally testing the use of our recently developed transition tempered metadynamics (TTMetaD) method to characterize membrane permeation of a relatively large molecule at the all atom level, and carefully comparing these results to those from coarse-grained MARTINI simulations. Following up on the work of Lopez et al. (17), we have herein used both CG and all atom molecular dynamics (MD) simulations employing TTMetaD to characterize the association of the toxin with DPPC membranes. We calculate membrane permeation free energy profiles for both isomers A and B, which exist in a 60:40 ratio. Our simulations show that, 1) the toxin has a strong preference for lipid phase with a 15-18 kcal/mol driving force to associate with a DPPC membrane, 2) the mechanism of permeation differs slightly for the A and B isomers with B preferring to associate with the glycerol groups, and 3) water facilitates permeation by associating with the polar tails of the toxin. Comparisons with the MARTINI simulations demonstrate how the CG approach gets the free energy of membrane association approximately correct, in concert with the parameterization procedure, but differs significantly in the mechanism of permeation and involvement of water. The insights gained are discussed in the context of how mycolactone-lipid interactions may influence the toxin’s association with and trafficking via membranes, as well as its distribution in the host environment. Not only will our continued understanding of this toxin-membrane dynamic be useful in development of diagnostics and adjunctive treatment approaches for Buruli ulcer disease, but it also serves as a fascinating model system of amphipathic-host interactions and lipid trafficking.

## METHODS

### All-atom Molecular Dynamics Simulation

In our all atom MD simulations, there are 50 × 2 DPPC molecules solvated with ~ 6300 water molecules (hydration ratio of 60), expanding to a roughly 5.75 nm × 5.75 nm region laterally (*x-y* directions), a size that is too large for a fully extended mycolactone (~ 3 nm) to interact with itself across periodic boundaries. The thickness of the simulation box (*z* direction) is about 10 nm. In order to compare our research with the previous work of Lopez et al. (17), we adopted the same MD simulation parameters for mycolactone, which were developed with the General Amber Force Field (GAFF) (22, 23) and restrained electrostatic potential (RESP) charges. To verify the legitimacy of these parameters, we compared them to those generated with the General Automated Atomic Model Parameterization protocol (24) and found them to be reasonably consistent. The DPPC molecules are modeled using an Amber-based force field (25) with corrections to balance the hydrophilic and hydrophobic forces (26). The water molecules are modeled using TIP3 force field (27). The initial structure of the DPPC lipid bilayer was prepared using CHARMM-GUI membrane builder (28) and equilibrated for 500 ns in water, and its thickness and area per headgroup were compared to the experimental value to confirm the bilayer had been equilibrated properly. One mycolactone molecule was then added into the bulk water in a random orientation. Thermostats (323K) using velocity rescaling with a stochastic term (29) were applied to the lipid bilayer and the rest of the system separately with a coupling time of 1 ps. A semiisotropic Berendsen barostat (30) (isotropic in the *x* and *y* directions, but independent in the *z* direction) was applied to control the pressure of the system with a coupling time of 5 ps. The cutoff distance for the short-range interaction list was 1.2 nm, which was updated every 20 steps. The long-range electrostatic interactions were computed using fast smooth Particle-Mesh Ewald (SPME) (31) and all hydrogen bonds were constrained by linear constraint solver (LINCS) (32). There were 5 replicas of 3.5 microseconds of TTMetaD simulations (integration time step: 0.2 fs) for both mycolactone A and B, with the initial velocities randomly sampled from a Boltzmann distribution. The all-atom MD simulations were carried out using GROMACS-5.0.4 (33) patched with a version of PLUMED2 (34) that was modified by the Voth research group to perform TTMetaD (publicly available in version 2.4 and beyond). In addition, all atom simulations of mycolactone B were run in the absence of any biasing force in order to verify the convergence of the role of hydration during permeation. The toxin was placed in the bulk solvent and allowed to associate with the membrane for a total of 160 ns of simulation.

### Transition-Tempered Metadynamics

Since membrane permeation is generally a slow process compared to the time scales accessible in MD simulations, enhanced sampling methods must be used. We have employed a variant of metadynamics (MetaD) (35-37), in which a bias energy (*V_G_*) is added to the Hamiltonian of the system along the simulation, through a small number of preselected collective variables (CVs) that represent the system of interest. The bias energy usually takes the form of the summation of Gaussians functions (Eq. 1), which are centered at the previously visited configuration in CV space, with a width of a tiny fraction of the CV scale.

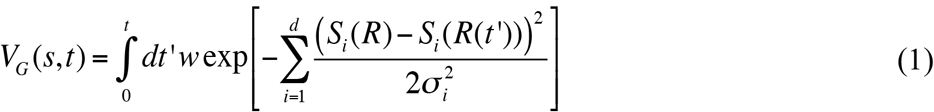

In the above equation, σ is the width of the Gaussian function and *w* stands for the bias energy incremental rate, i.e. the height of the Gaussian functions (*w_0_*) divided by its deposition stride (*τ*). *S_i_*(*R*(*t’*)) is CV value evaluated from the configuration of the system at time *t’* with the index *i* representing the *i*^th^ CV. As a result of the bias energy, the system is discouraged from revisiting the previously visited point, pushing it away from local energy minima such that it can explore higher-energy regions in CV space. Eventually, the motion of the system in CV space becomes diffusive and ergodic, indicating the bias energy has offset the “resistance” of the system. As a result, the underlying free energy of the system can be estimated as the negative of the MetaD bias energy. This approach, originally introduced in 2002 (35), is often regarded as non-tempered MetaD today because the height of the Gaussian in Eq. 1 remains constant throughout the simulation. Although non-tempered MetaD is generally effective in exploring CV space, the bias energy oscillates around the true underlying potential of the mean force (PMF) instead of converging asymptotically, in contrast to the fact that the PMF is a time independent property. Calculating time average of the bias energy from non-tempered MetaD (only after the motion of the system becoming diffusive) has been proposed as a solution to this convergence issue, but practically, we have found that this either shows false convergence or destabilizes the simulation due to too much bias energy being added (38, 39).

The incremental bias additions (i.e. *w* in Eq. 1, depending on the height of individual Gaussian, *w_0_*) need to zero to achieve the true convergence of the bias energy. Well-tempered metadynamics (WTMetaD) (40) is such method that replaces the time-independent incremental bias additions (*w*) with a time-dependent quantity, *w*(*t*):

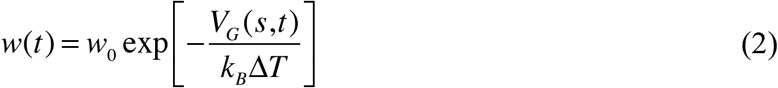

The bias energy incremental rate decreases exponentially with respect to the local bias energy. The parameter *ΔT* tunes the amount of decrement and is chosen prior to the simulation. In principle, WTMetaD has been proven to converge the bias energy asymptotically to a linearly scaled inverse of the underlying free energy (41). Despite the success of WTMetaD, it presents a trade-off between the exploration and the converging in WTMetaD: a fast-growing bias potential (with large *ΔT*) leads to a noisy bias energy, hindering the convergence; a slow-growing (with small *ΔT*) leads to a smooth bias potential, hindering the exploration. It has been suggested that *ΔT* should be chosen according to the largest free energy barrier so that (*ΔT* + *T*)*k_B_* has the same order of magnitude as the barrier (40). However, the barrier height is usually the most interesting feature of the system and often unavailable, therefore the efficiency of WTMetaD can sometimes be limited considerably.

Converging like WTMetaD, transition-tempered metadynamics (TTMetaD) (42) overcoming the aforementioned trade-off by tempering the bias energy incremental rate with respect to overall progress of the MetaD sampling. Namely, a global property *V^*^*, the minimal bias on the maximally biased path among all the continuous paths *s*(*λ*) connecting all of the preselected points in the CV space, replaces the local bias energy *V_G_* (Eq. 2):

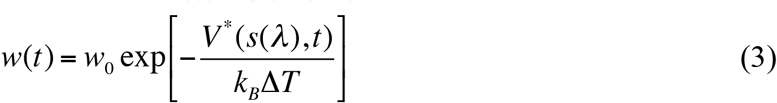

In TTMetaD, the preselected points are the basins on the underlying free energy surface, separated by a significant barrier. Intuitively, *V^*^* can be understood as the amount of the bias energy needed for the least-likely point in the CV space to be sampled, usually corresponding to the transition state (TS) region. The *V^*^* remains zero before the basins are connected through TS, thus TTMetaD explores with a full Gaussian. Once the TS region has been sampled, TTMetaD can afford a more aggressive tempering (small *ΔT*) of the bias energy incremental rate to converge the simulation. TTMetaD has demonstrated its advantages of converging rapidly and asymptotically in various studies of complicated biophysical systems (38, 39). The 2D PMF is estimated from the reverse of the average of the bias energy from five independent replicas, and then diagonally symmetrized with respect to the center of the membrane and toxin orientation. This method has previously demonstrated its efficiency in converging TTMetaD simulations, especially for the lipid bilayer permeation. The minimum free energy path (MFEP) on the 2D PMF (black curve in Figure 4), which represents the most common pathway in a large ensemble of the permeation processes, was calculated with a zero temperature string method (43). 1D PMF is directly obtained by using the average MFEP from five independent replicas (44).

### Coarse Grained Molecular Dynamics Simulation

In principle, using a CG approach like MARTINI (45) could enable simulations of mycolactone interacting with membranes at much larger length scales and longer time scales. Thus, our intention in comparing our all atom MD results with those from MARTINI is to verify its reliability in modeling this complicated toxin-membrane interaction. To make as clear of a comparison as possible, the same methods were employed and the same system size was used as those in the all atom MD simulations (adjusting for the MARTINI CG water bead representing 4 water molecules). We focused the comparison on the permeation of mycolactone B only. The MARTINI CG force field parameters were adapted from Lopez et al. (17), where the MARTINI CG representation of mycolactone B was obtained from the *insane* protocol (46). After equilibration, the dimension of the MARTINI CG simulation box was about 5.5 nm × 5.5 nm × 6.6 nm. Although the length along the z direction decreased significantly from the all-atom scale, this was still deemed sufficient for the characterization of permeation. The thermostat, barostat, and long-range electrostatic interactions were kept consistent with those used in the all-atom MD simulations. Various short-range interaction cutoff distances and neighbor list update frequencies were tested, and final values of 1.4 nm/updating every 10 steps were chosen for the stability of the simulation. These are different from those previously used (17). There were 6 replicas of 20 microseconds (dummy integration time step: 0.2 fs) for the TTMetaD MARTINI CG simulations.

### Collective Variables

In MetaD, the bias energy is deposited into CV space to accelerate exploration of the free energy landscape. In general, the CV needs to capture the process of interest (e.g., a reaction coordinate for a chemical reaction), delineating between the starting and ending states (e.g., reactant and product). Ideally, it should also capture the slow degrees of freedom that limit progression from the start to end state. Using TTMetaD we have studied the permeation of small organic molecules though lipid bilayers, and found that at least two CVs are often necessary – one that describes the center of mass translation of the permeant through the membrane and another that describes molecular reorientation during permeation. Since mycolactone is a relatively large and flexible molecule, both the center of mass of the whole molecule and the center of mass of the lactone ring were tested as CVs to define the translation of the permeant; the latter was chosen because it demonstrated faster convergence while the former failed to delineate between a wide range of configurations with slow interconversion, thereby limiting convergence. The orientation CV was defined as the angle between the vector connecting the hydroxyl groups on the end of the northern and southern tails, and the vector normal to the lipid bilayer. An illustration of the two CVs is shown in Figure 1. The same CVs are defined for the MARTINI CG simulations through the corresponding beads.

**Figure 1.**
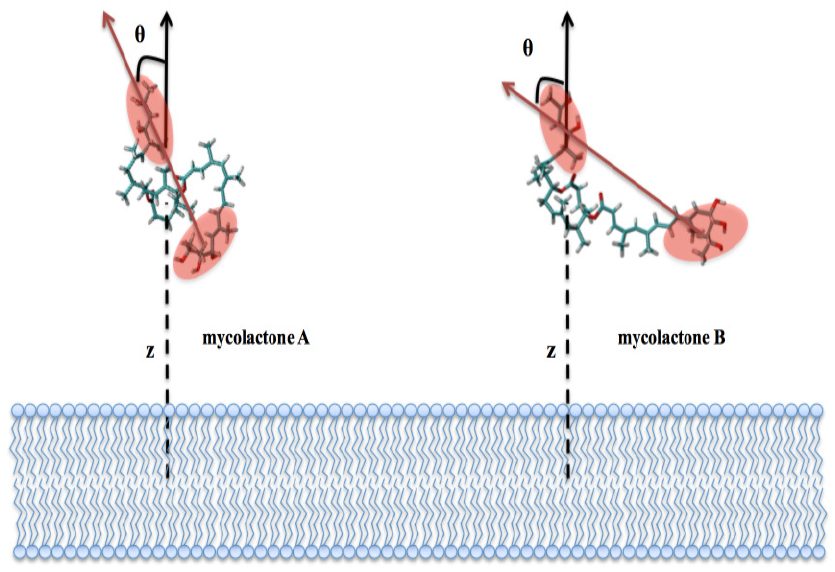
Definition of CVs for mycolactone A/B permeation through a DPPC lipid bilayer.

## RESULTS AND DISCUSSION

### Interconversion between Mycolactone A and B

Mycolactone is a polyketide composed of an invariant 12-membered lactone ring with two highly unsaturated acyl side chains (see top panel of Figure 2). The shorter side chain (often called the ‘Northern’ chain) is invariant, whereas the longer (‘Southern’) chain varies in different congeners of mycolactone. The most cytotoxic and common congener is mycolactone A/B (*in vitro*), which exists in two isomeric forms in a ratio of 60:40 (A:B) under common laboratory conditions and light. We sought to perform our all-atom simulations on both isomers to see if, within the limited accuracy of the force field and simulation techniques, the permeation process differs. We first considered interconversion between the two isomeric forms. One mycolactone B molecule solvated in 4400 water molecules was simulated for 1.6 microseconds.

**Figure 2.**
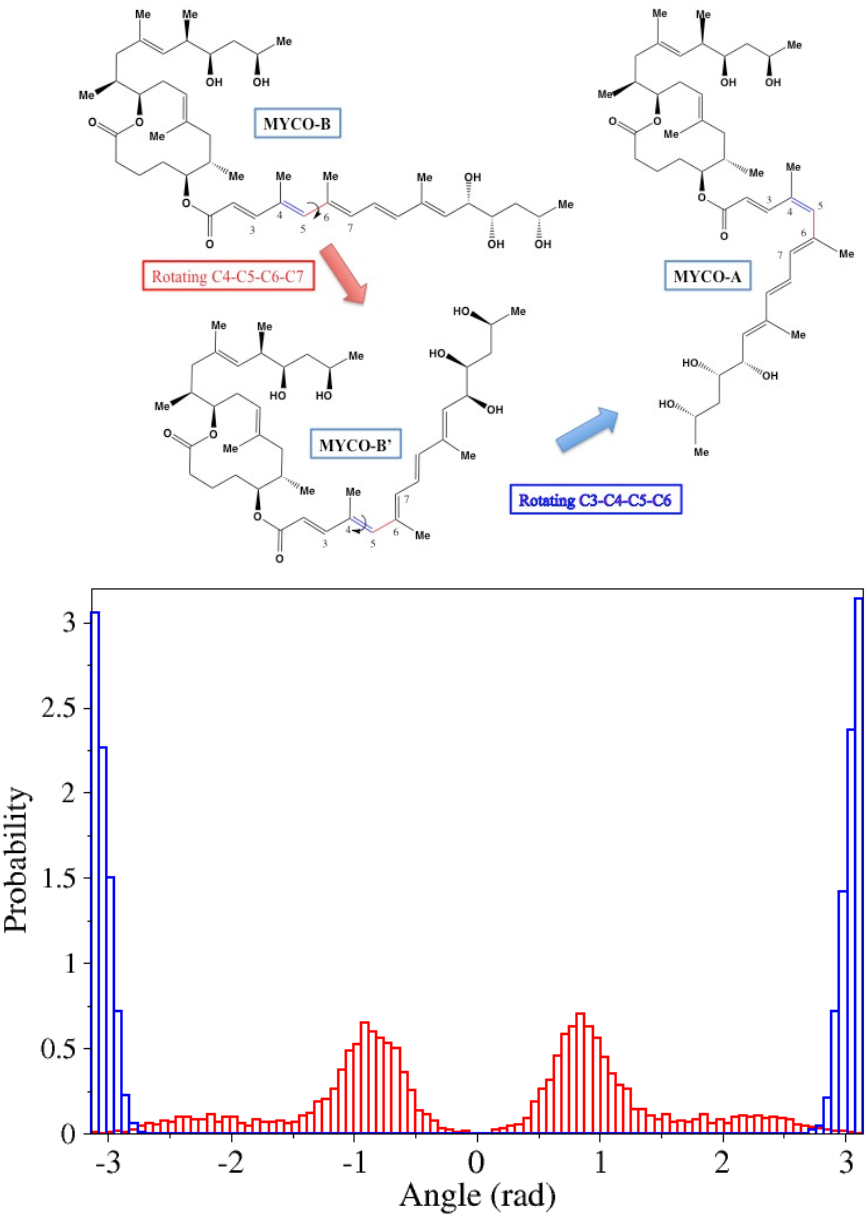
Top panel: The conformation of mycolactone A and B. Bottom panel: The normalized distribution of dihedral angle 3-4-5-6 (blue) and 4-5-6-7 (red) from mycolactone B and water simulation.

Although the dihedral angle of the single bond (4-5-6-7) is flexible, leading to the conformational change between mycolactone B and B’ (Figure 2); the 3-4-5-6 (double bond) dihedral angle distribution from the simulation shows it is very stable, as expected, on a microsecond timescale. Therefore, the permeation of mycolactone A and B are modeled separately in atomistic simulations in this manuscript.

### Permeation PMF from All-atom Simulations

The DPPC membrane and mycolactone molecule (in all atom and CG representation) are depicted in Figure 3. The 2D potential of mean force (PMF) of the permeation of mycolactone B through DPPC lipid bilayer is shown in Figure 4, with the inserted snapshots illustrating representative configurations. Our results confirm the strong affinity between mycolactone and lipid bilayers, reported by both experiments and MARTINI CG simulations (14, 17).

**Figure 3.**
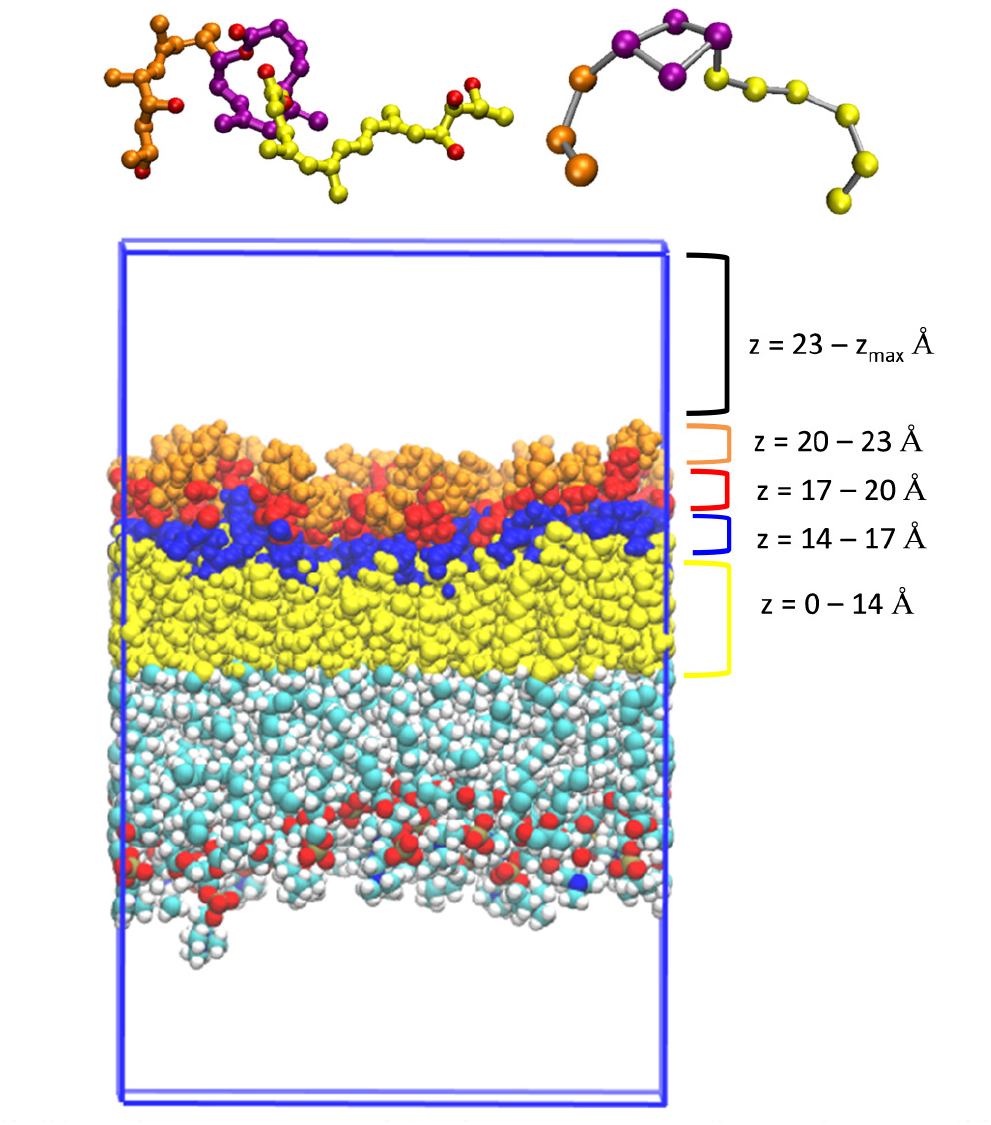
Top panel: All-atom (left) and coarse-grained (right) structures of mycolactone. The northern tail, southern tail and lactone ring are orange, yellow and purple, respectively. Oxygen atoms of mycolactone in all-atom structure are red. Bottom panel: The DPPC membrane structure from all-atom simulations. The upper monolayer is colored based on different lipid regions: hydrophobic tail (yellow), glycerol (blue), phosphate (red) and choline (orange).

**Figure 4.**
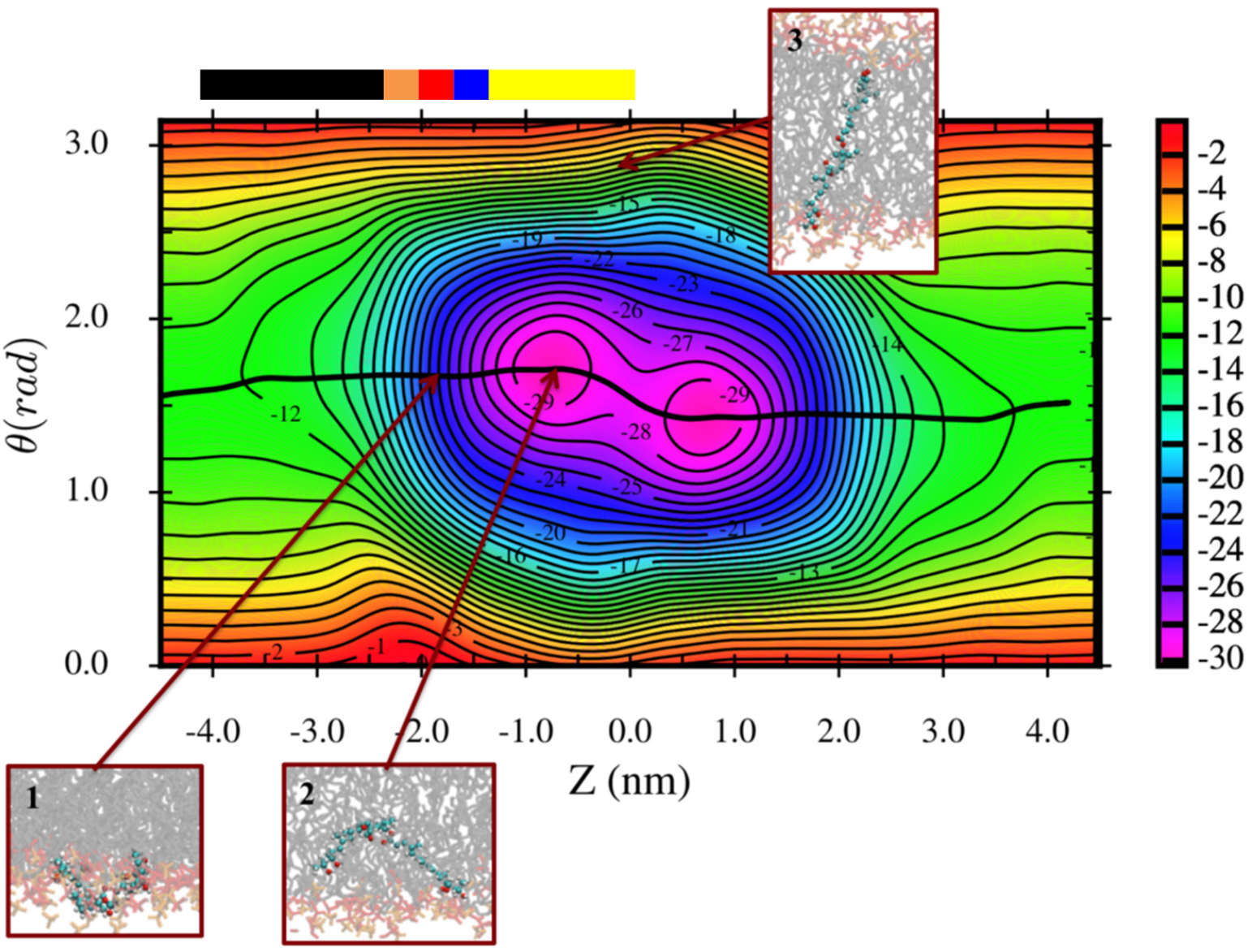
2D PMF of mycolactone B permeation through DPPC lipid bilayer. The black line represents the MFEP and the inserted figures depict the representative configurations. The toxin is colored by atom type while the DPPC head groups, glycerols, and tails are colored with orange, red, and grey, respectively. The membrane spans −2.0 < Z < 2.0 nm. The color bar corresponds to the colored regions of the DPPC membrane shown in Figure 3.

The MFEP demonstrates how the permeation process is dominated by hydrophobic-hydrophilic interactions between mycolactone and DPPC lipids. When mycolactone approaches the lipid bilayer, the two hydrophilic tail ends (including two and three hydroxyl groups on the northern and southern tails, respectively) interact with the hydrophilic head groups of the lipid bilayer (configuration #1, Figure 4). This mycolactone-tail/DPPC-head interaction is strongly favored in free energy and remains stable as the toxin moves into (and out of) the bilayer; therefore, the orientation CV value (*θ*) does not vary significantly from −4 nm to −1 nm (and 1 nm to 4 nm). As the permeation continues, the hydrophobic lactone ring submerges into the tail groups of the lipid bilayer (configuration #2, Figure 4). This configuration corresponds to the lowest free energy on the 2D PMF. To permeate to the other side, the toxin flips, keeping hydrophobic lactone ring in the lipid tail region and swapping the hydrophilic toxin tails to interact with the head groups on the other lipid leaflet. Finally, the lactone ring is pulled from the tail group region to finish the permeation process. The preference of the toxin tails to interact with the polar head groups is also apparent in the 1D free energy profile of the MFEP (Figure 6), which shows a slight increase in the free energy in the middle of the membrane (Z = 0.7 to 0 nm). It is important to point out that the stretched configuration of mycolactone, where the northern and southern tails interact with opposing leaflets (configuration #3, Figure 4), corresponds to a high free energy, disfavored configuration. This is in contrast to the previous MARTINI simulations showing the extended configuration as a favored metastable minimum, and suggesting the permeation process involved such an extension as the toxin passes from one side of the membrane to the other (17).

The permeation of mycolactone A through the DPPC bilayer shows a very similar PMF (Figures 5 and 6) as the permeation of mycolactone B, indicating a similar mechanism. A small but interesting difference is the lack of change in the orientation CV value (*θ*) in the 2D PMF (Figure 5) during the flip of the toxin from one leaflet to the other. This is also apparent in the 1D free energy profile (Figure 6) which is flat in the middle of the membrane as opposed to showing a slight free energy increase as mycolactone B does. This can be attributed to a structural difference. With the rotation of a single bond, the two hydrophilic tails in mycolactone A can flip around to hydrogen bond to each other or to collectively coordinate water molecules. This better satisfies the polar tail interactions making it easier for the toxin to pass into the lipid tail region. In addition, mycolactone A pulls water molecules deeper into the lipid bilayer (close to the middle of the membrane) than mycolactone B does, as described in the next section. Similar to mycolactone B, the stretched configuration remains unfavorable for mycolactone A (e.g. configuration #3, Figure 5).

**Figure 5.**
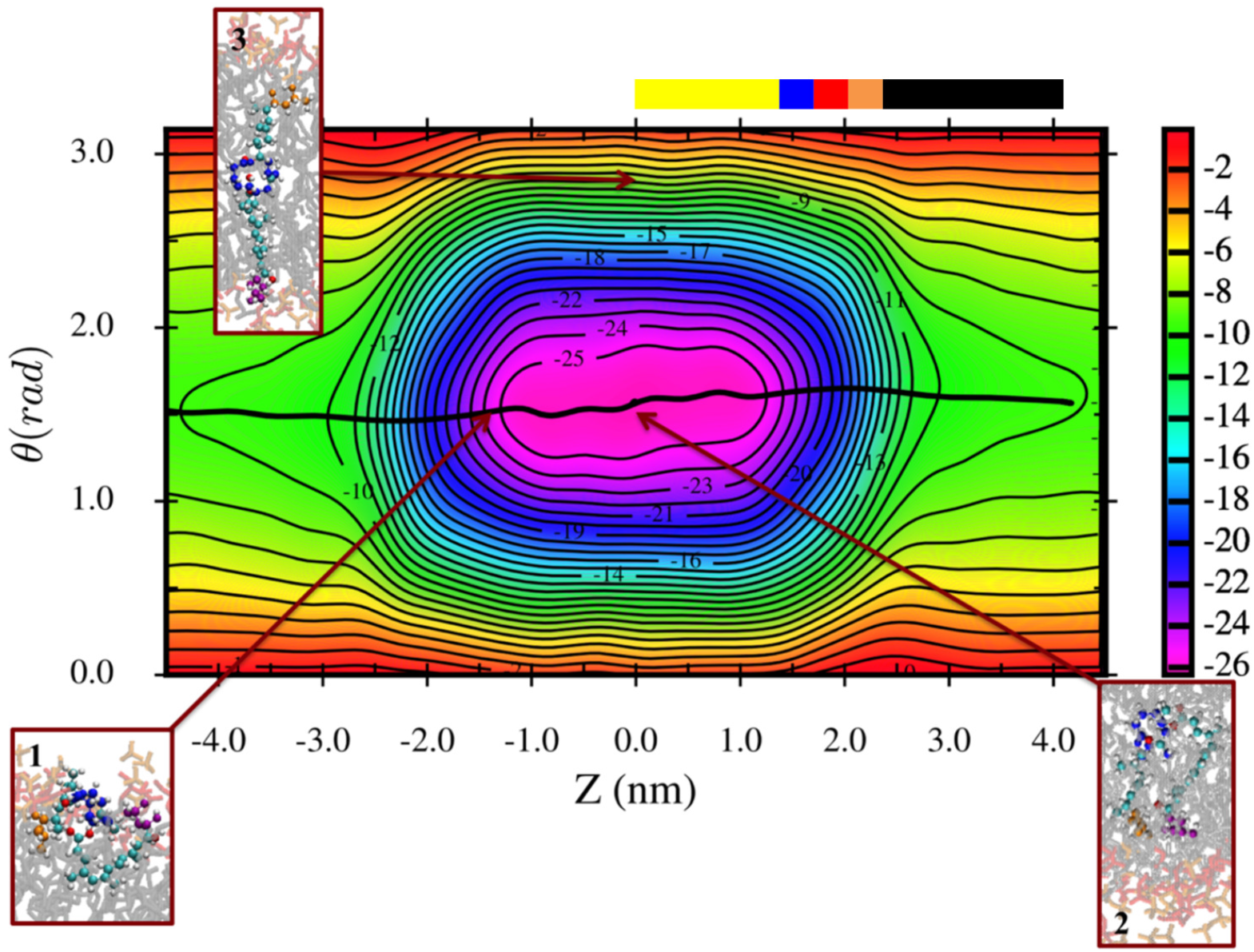
2D PMF of mycolactone A permeation through DPPC lipid bilayer. The black line represents the MFEP and the inserted figures depict the representative configurations. The northern tail, southern tail, and lactone ring are colored with orange, purple, and blue, respectively. The DPPC head groups, glycerols, and tails are colored with orange, red, and grey, respectively. The color bar corresponds to the colored regions of the DPPC membrane shown in Figure 3.

**Figure 6.**
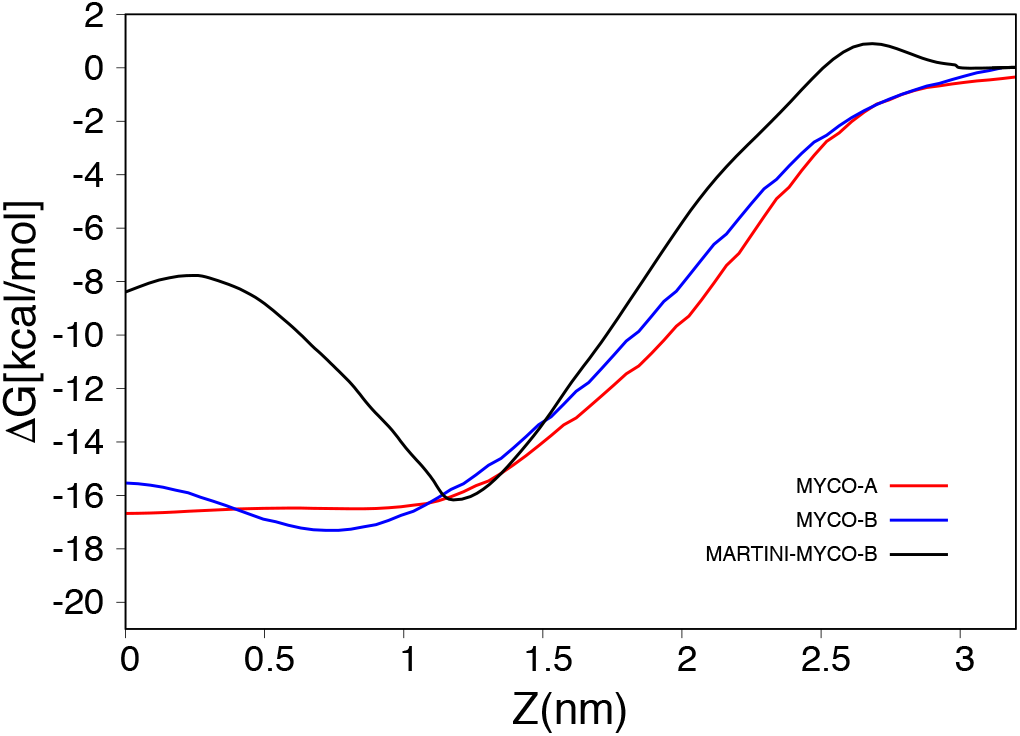
1D free energy profiles of the permeation of mycolactone B (atomistic and MARTINI CG) and mycolactone A (atomistic) with respect to the insertion depth of the lactone ring into the lipid bilayer.Error bars (0.2-3 kcal/mol) not shown for clarity.

### The Role of Hydration in Mycolactone Membrane Permeation

It is intriguing to focus on the role of water during the permeation of mycolactone. As shown in the permeation PMFs of both isomers, the most favored (lowest in free energy) configurations minimize the hydrophobic-hydrophilic interactions – the highly hydrophobic lactone ring is buried in the hydrophobic lipid tails, and the hydrophilic tails are stretching out to the hydrophilic lipid head groups and/or interacting with water molecules. We first analyzed the number of water molecules interacting with mycolactone as a function of the z distance of the lactone ring to the membrane center in the unbiased (mycolactone B) and biased all-atom (mycolactone A and B) simulations, as well as the biased MARTINI CG (mycolactone B) simulations. A water molecule is considered to be interacting with mycolactone if the COM distance between the oxygen atom of water and any oxygen atom of mycolactone is less than or equal to 3.04 Å for all-atom simulations. In concert, a water bead is considered to be interacting with mycolactone if the COM distance between the water bead and any bead of mycolactone is less than or equal to 4.9 Å for MARTINI CG simulations. The probability distributions of number of water molecules interacting with mycolactone A and B for biased and unbiased all-atom simulations are all in good agreement (see Figure 7). Mycolactone continues to interact with water molecules when the lactone ring is near the center of membrane (z = 0). However, there is a striking difference between the hydration profiles of all-atom and MARTINI CG simulations. Contrary to the all-atom simulations, the MARTINI mycolactone has a very low probability of interacting with a water bead when the lactone ring moves to through center of membrane.

**Figure 7.**
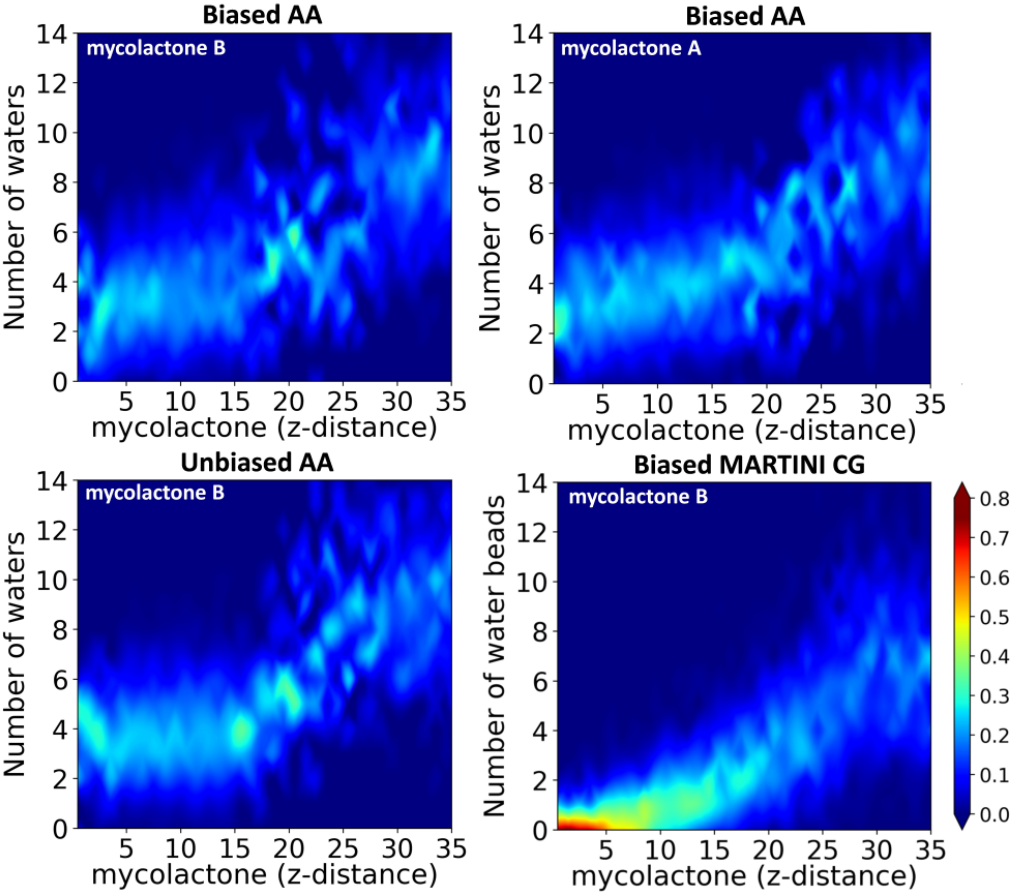
Probability distributions of number of water molecules interacting with mycolactone with respect to the insertion depth of the lactone ring into the lipid bilayer from unbiased and biased all-atom, and biased MARTINI coarse-grained simulations.

To better understand *where* these coordinating waters were in the membrane, we next analyzed the position distributions of coordinating water molecules as a function of the z position of the lactone ring. The position distribution of water molecules is obtained by calculating the z-distance between each water molecule interacting with mycolactone and the center of membrane (Figure 8). Similar to the number distributions of water molecules, the position distributions in unbiased and biased all-atom simulations of mycolactone B are reasonably consistent with each other. The distribution of the unbiased all-atom simulation is not uniform because the toxin quickly penetrates into the membrane and spends most of the time inside the lipid bilayer. In addition, mycolactone does not spend much time in the center of the membrane, causing sparse sampling, in unbiased simulations due to its preference to maintain tail-head group interactions (i.e., due to the minimum at Z = 0.7 nm). An interesting subtle difference is that the biased simulations have a wider distribution of water positions, mostly associating with water further away from and to a lesser degree pulling water further into the center of membrane, as compared to the unbiased simulation. This suggests that hydration itself is a slow degree of freedom that is not exhaustively sampled in the TTMetaD simulations. However, the difference is so slight that explicit sampling of a hydration CV is not expected to change the free energy profile significantly. Importantly, both the unbiased and biased all-atom simulations demonstrate deep penetration of water molecules with the insertion of mycolactone into the membrane. Thus, the penetrating water molecules play an important role in the permeation process – compensating for lost hydrophilic interactions. In comparison, mycolactone A pulls water molecules more deeply into the lipid tail region (close to the membrane center – see Figure 8 lower panel), suggesting a more compact structure during permeation. The representative configurations of each system shown in Figure 9 capture the differences in water coordination and structure between the two isomers. The ability of mycolactone A to pull water molecules due to its compact structure can be also seen in Movie S1 in the Supporting Material. In contrast, the position distribution of water beads in the biased MARTINI CG simulations is quite different (middle panel of Figure 8 and Figure 9). When the lactone ring is near the center of membrane (z = 0), the water beads are around or just below the lipid head group region (z = 10 – 20 Å) revealing no deep penetration of MARTINI water beads. As discussed below, this is a likely contribution to the free energy barrier close to the membrane center in the MARTINI PMF.

**Figure 8.**
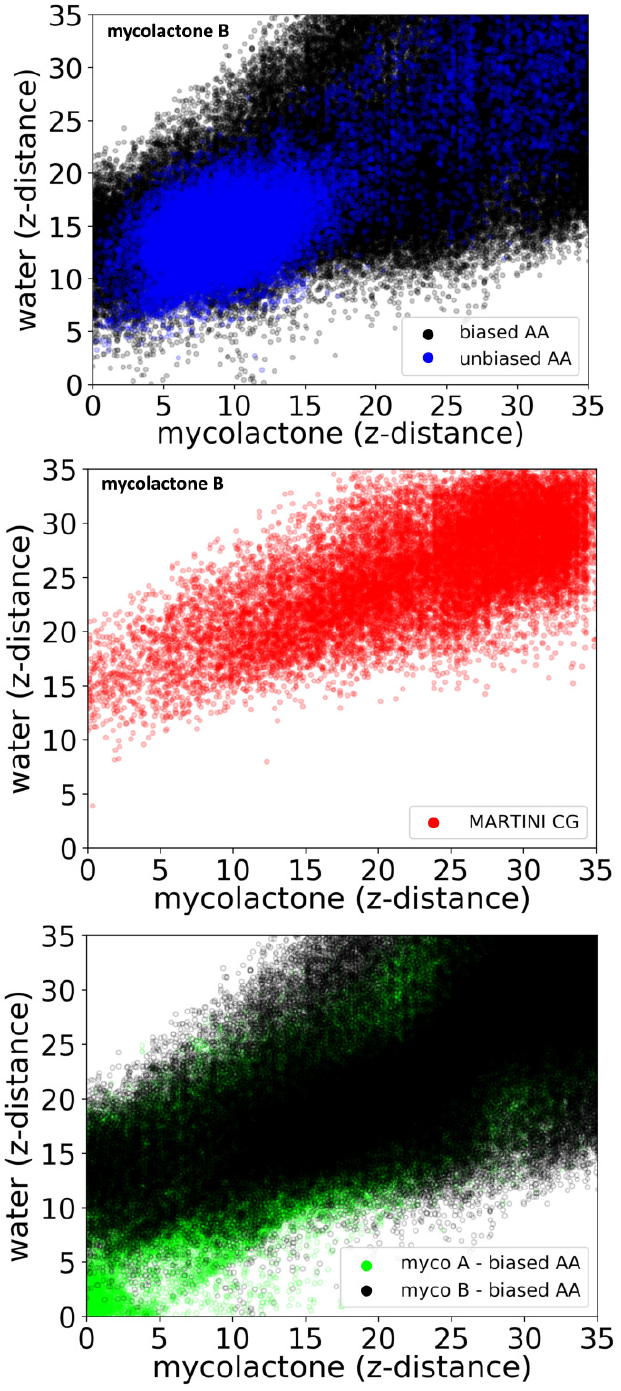
Position distributions of water molecules/beads interacting with mycolactone with respect to the insertion depth of the lactone ring from unbiased and biased all-atom, and biased MARTINI CG simulations.

**Figure 9.**
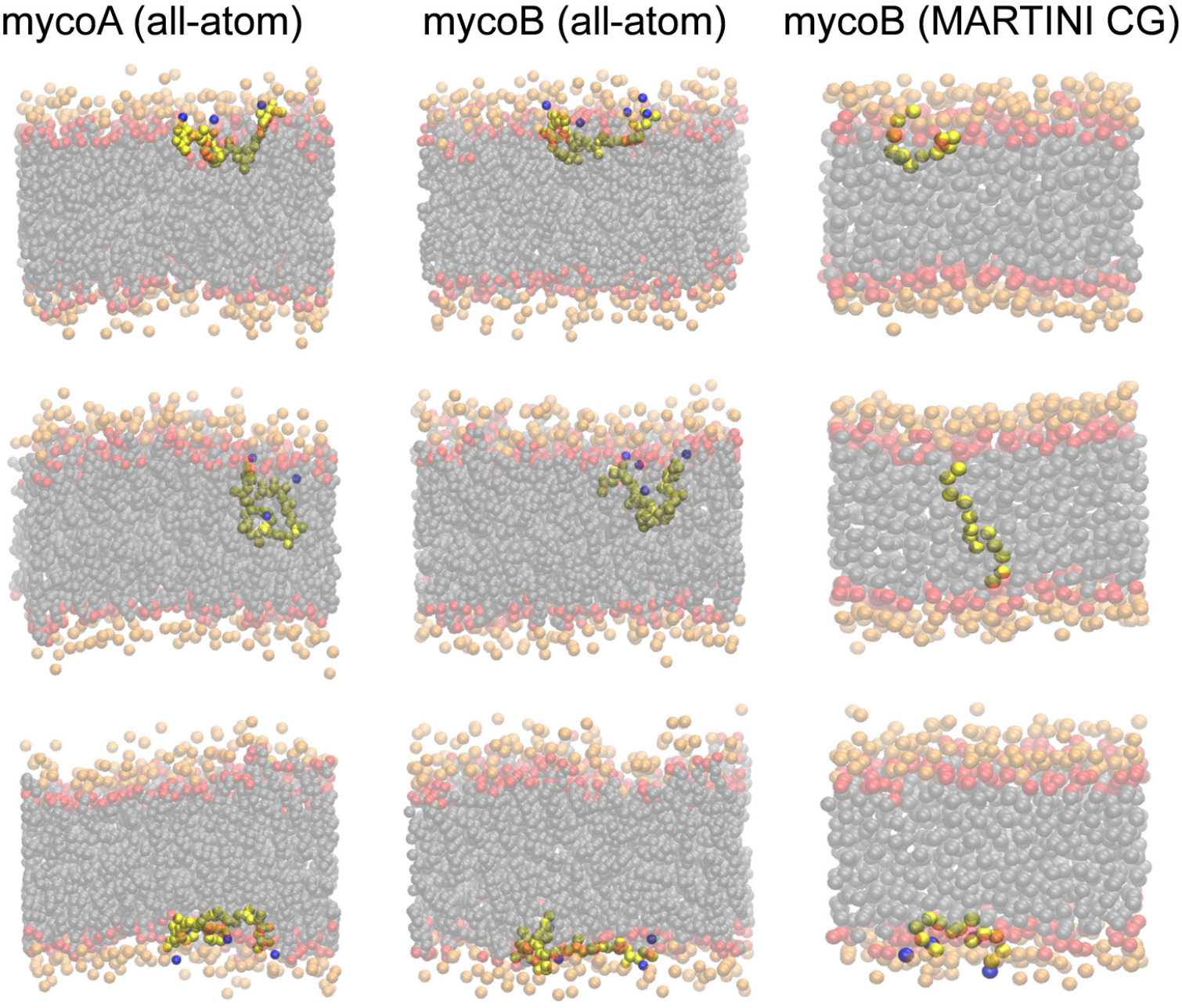
The snapshots depict the representative configurations from all-atom simulations of mycolactone A and B (the left and middle columns), and from MARTINI CG simulations of mycolactone B (the right column). The mycolactone and water molecules are colored with yellow, and blue, respectively. The DPPC head groups, glycerols, and tails are colored with orange, red, and grey, respectively.

### Comparison with the MARTINI CG Simulation

Coarse grained simulations can extend the accessible temporal and spatial scale of molecular modeling, which is sometimes essential to connect with the biological process of interest.

However, the accuracy of CG models, especially for molecules they were not originally parameterized for (e.g., organic molecules like mycolactone), is often unknown. In order to confirm a CG simulation is trust worthy, it is generally essential to verify the accuracy of a coarse-grained model by comparing it to more detailed models and experimental data. In addition, such comparisons can be useful in improving CG methods. Herein, we compare the permeation PMFs and mechanisms of mycolactone B through DPPC lipid bilayers from MARTINI CG and all-atom simulations.

As discussed in the methodology section, the MARTINI CG system was directly coarse grained from the all-atom system, consistent with a previous publication (17). In contrast to that work, the methods used herein are consistent with the methods used for our all atom simulations to provide as robust of a comparison as possible. Transition-tempered metadynamics was employed to calculate the permeation PMF with exactly the same CVs (after coarse graining) as in Figure 1. The converged 2D PMF is depicted in Figure 10, and the 1D free energy profile is in Figure 6. Compared to the previously published PMF, our PMF shows differences in shape (less stabilization at Z=0 and smoother transitions) and thermodynamics (*Δ G* for membrane association of −16 kcal/mol herein vs −12 kcal/mol previously), which we attribute to treatment of cut off interactions and methods used to sample the PMF.

**Figure 10.**
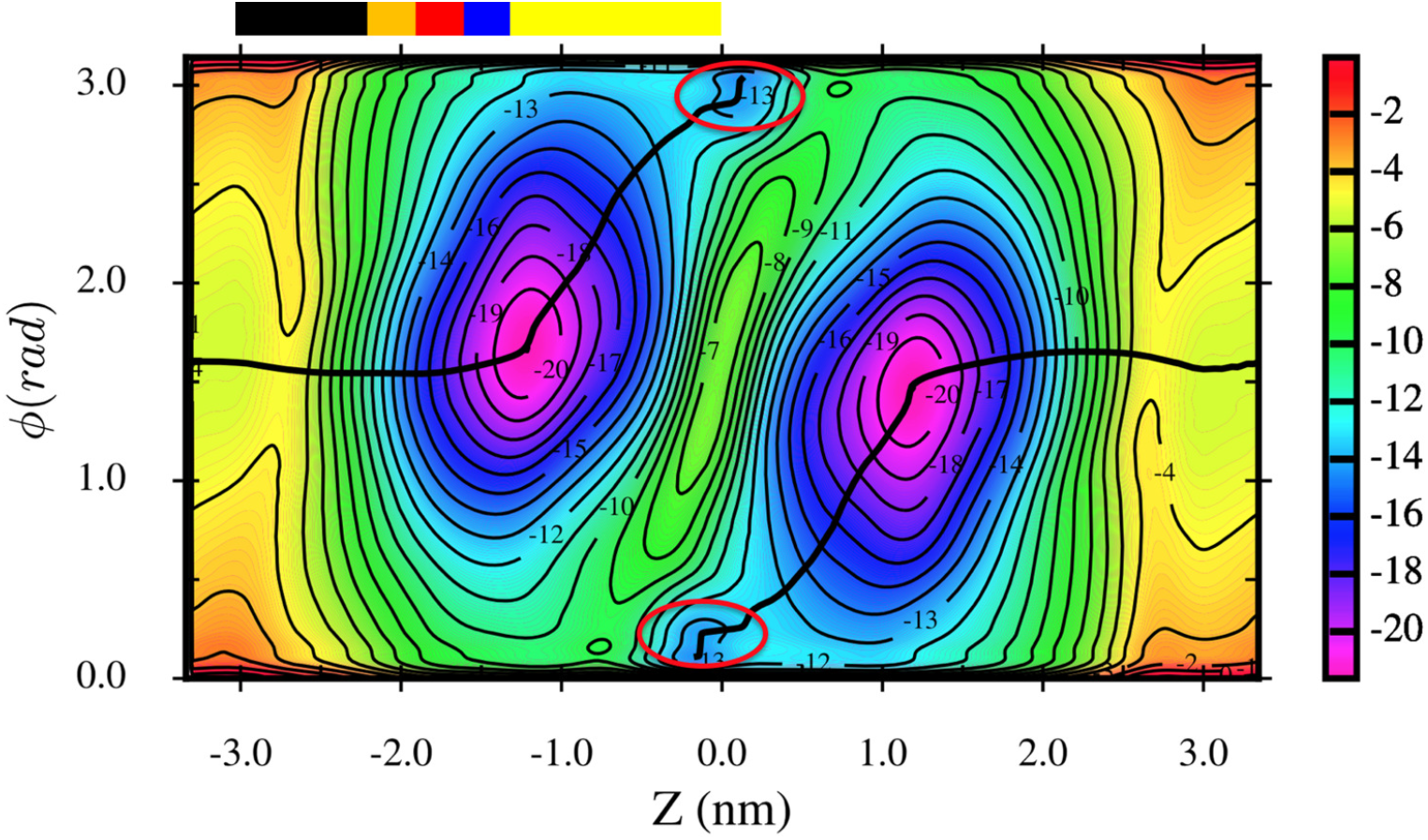
2D PMF of mycolactone B permeation through DPPC lipid bilayer from MARTINI CG simulations. The black line represents the MFEP. The red circles illustrate the configurations where mycolactone is full stretched to head groups of both leaflets, corresponding to configuration #3 in Figures 3 and 4. The color bar corresponds to the colored regions of DPPC membrane given in Figure 3.

Comparing our all-atom simulations to our MARTINI simulations we find that the CG model also predicts a strong driving force for membrane association and affinity for the toxin tails to interact with the lipid head groups, though this interaction localizes at Z = 1.3 nm as opposed to 0.7 nm as it did in the all atom simulations (Figure 6). Again, this optimizes the hydrophobic-hydrophilic interactions. However, the differences are very clear: the MARTINI CG model predicts a barrier for the toxin to pass through the lipid tail region and a metastable fully stretched configuration (marked with red circles in Figure 10), where the two tails of the mycolactone molecule interact with the head groups on opposing leaflets. This configuration is strongly disfavored according to the all-atom simulation (configuration #3 in Figure 4). It is encouraging that the membrane association energies are very similar from MARTINI CG (~ 17 kcal/mol) compared to all-atom simulations (~ 18 kcal/mol). However, the large barrier for dissociation of the toxin tails from the lipid head groups (an increase in free energy of ~9 kcal/mol going from Z = 1.3 nm to 0.3 nm) and the metastable extended configuration demonstrate a very different mechanism of permeation. It is important for the continued use and development of CG models to understand the origin of these differences. An obvious contributing factor is the approximate nature of the CG parameterization and thus differences in the CG and all atom toxin-lipid interactions. In addition, however, we believe the role of water molecules could play an important role. As discussed in the previous section and shown in Figure 8, there is no deep water penetration into the lipid tail region in MARTINI CG simulations, in contrast to the considerable penetration of water molecules observed in unbiased and biased all-atom simulations. These penetrating water molecules stabilize the toxin in the lipid tail region, enabling a balance between hydrophilic and hydrophobic interactions in the membrane environment. This not only helps to eliminate the barrier observed for permeation in the CG simulations, it also alters the mechanism by which the toxin permeates. In the CG simulations, the cost of transitioning to the extended conformation, which maintains polar interactions with the head group beads on either side of the membrane, is outweighed by the cost of losing polar interactions in the tail region. Other factors, such as the overly rigid bending modulus and increased lipid diffusivity of MARTINI membranes could additionally be contributing factors to the different PMFs and permeation mechanisms (47-49).

## CONCLUSIONS

Mycolactone is a complex and multifunctional molecule that targets various structures in the cell and likely influences membrane microdomains. This cytotoxic macrolide is known to play the central role in the progression of Buruli Ulcer disease by disrupting multiple cellular functions. Effective diagnostic methods are still missing, partially due to the association of mycolactone with lipidic structures, enabling this lipid-like molecule to hide from tracking agents. Moreover, the association of mycolactone with lipid-carrying moieties could explain how the toxin evades the secondary immune response and travels far from the site of infection. Therefore, understanding the toxin’s interactions with lipids, permeation mechanism and localization is important to understand its pathogenesis and to design effective diagnostics. Furthermore, the findings from mycolactone can be extended to other amphipathic molecules to understand host-pathogen interactions in general. Towards this aim, we have presented all-atom and MARTINI CG TTMetaD simulations that characterize the behavior of two isomers (mycolactone A and B) interacting with a DPPC membrane.

Our all-atom simulations demonstrate a strong association between toxin and the membrane with a free energy of ~17 kcal/mol. They also confirm the expected role of hydrophobic-hydrophilic interactions for any amphiphile, with the macrolide ring burying in the lipid tails and the polar toxin tails interacting with lipid head groups and water. There is a slight but remarkable difference in the mechanism of permeation between two isomers. Only mycolactone B exhibits a shift in the orientation and a small free energy increase when it flips from one leaflet to the other (Figure 4 vs Figure 5, and Figure 6). This variation can be explained by the structural difference between two isomers: the two tails of mycolactone A can hydrogen bond with each other and collectively coordinate water molecules, allowing it to cross the lipid tail region more easily. Our all-atom simulations also demonstrate that water molecules play a critical role in the permeation process. Both isomers are stabilized by coordination to ~2-4 water molecules during permeation.

Although the free energy driving force for membrane insertion is quite similar in the MARTINI CG simulations, the mechanism of permeation of mycolactone B differs substantially from our all atoms results. The fully stretched configuration is strongly disfavored by both isomers in all-atom simulations, but is favored in the MARTINI CG simulations. This discrepancy is consistent with previous publication showing that the extended configuration plays a dominate role in the MARTINI-based permeation mechanism (17). We also observe quite different behavior of toxin hydration in the MARTINI CG simulations: water coordination is disfavored during permeation and there is virtually no deep-water penetration into the MARTINI membrane. Since water molecules penetrating into the lipid tail region will alter the balance between the hydrophilic and hydrophobic interactions, this is expected to contribute to the altered free energy profile and mechanism of permeation.

While our findings are specific to one lipidic component in membranes, DPPC, this work represents an important step forward in understanding the lipid-specific interactions that undoubtedly play an important role in how mycolactone penetrates host cells, rapidly travels to the endoplasmic reticulum, and subsequently travels far from the site of infection in the host vasculature system. They will also play a defining role in the toxin’s expected inhibition of membrane microdomain formation. How the toxin interacts with different lipids and how it localizes to different membrane regions and leaflets will strongly influence these processes. These are the processes we seek to understand. Given the mechanistic discrepancies presented herein, it is unlikely that the MARTINI based CG models will be capable of delineating lipid-specific interactions with fidelity. Thus, this work points to the importance of using all atom simulations to further probe lipid trafficking processes where lipid-specific interactions likely play a role. It also suggests ways in which MARTINI based CG models could be improved – focusing on water-enriched membrane interactions.

## AUTHOR CONTRIBUTIONS

J.M.J.S. designed the research. R.S. performed the simulations. F.A., R.S. and J.M.J.S performed the analysis; and all authors contributed to writing the paper.

## ACKNOWLEDGMENTS

This work was supported by the National Institute of Allergy and Infectious Diseases, R01-AI113266. Simulations were performed using resources provided by the University of Chicago Research Computing Center (RCC).

